# Satellite Glial Cells Drive Homeostatic Synaptic Structural Plasticity in Sympathetic Neurons

**DOI:** 10.64898/2026.05.10.723591

**Authors:** Joshua Harrison, Ellie Greene, Aaron Yang, Jumana Akoad, Laurie Chen, Rui Gong, Xianghan Liu, Susan Birren

## Abstract

Sympathetic neuronal (SN) activity critically regulates the development and function of peripheral organs and tissues. The demonstration of activity-dependent modulation of SN output suggests that compensatory forms of plasticity could contribute to maintaining the stability of sympathetic circuits. Such plasticity mechanisms could act to restrain SN hyperactivity, a key driver of hypertension in humans and in the spontaneously hypertensive rat (SHR). In this study we examined how long-term changes in activity impact synaptic properties in postnatal sympathetic neuron cultures using chemogenetic and pharmacological manipulations and by examining the effects of enhanced activity of SHR neurons. We showed that bidirectional changes in neuronal activity resulted in homeostatic shifts in synaptic density to counteract long-term activity manipulations. In the absence of sympathetic satellite glial cells (SGCs) there was no synaptic compensation in the cultures. Direct chemogenetic activation of SGCs was sufficient to drive a decrease in synaptic sites and neuronal activity, while glial inhibition blocked activity-dependent synaptic compensation, demonstrating a role for the SGCs in homeostatic regulation of synaptic properties. We found that the SGCs responded to cholinergic signaling by downregulating the expression of the synaptic regulators NGF and TNFα, suggesting that reciprocal signaling between SNs and SGCs acts to stabilize sympathetic output during long-term changes in circuit activity. Finally, we showed that these plasticity mechanisms are disrupted in postnatal SHR neurons, with an attenuated neuronal response to glia signaling during synapse formation and activity-dependent plasticity. Taken together, this work describes a new homeostatic activity-dependent plasticity mechanism in the peripheral nervous system.

## Introduction

Activity-dependent plasticity describes a set of mechanisms for adjusting neuronal synaptic and firing properties in response to changes in network activity. These mechanisms drive both feed-forward and homeostatic changes that interact to fine-tune synaptic function and intrinsic neuronal properties (Sweatt, 2016), allowing for bidirectional constraint of firing rates within a preferred range (Wen and Turrigiano, 2024). Numerous studies highlight the critical role of this process in shaping brain circuits (Grienberger and Magee, 2022; Bissen et al., 2025; Le et al., 2025). In the peripheral sympathetic nervous system, tetanic stimulation of preganglionic inputs to the sympathetic ganglion strengthens synaptic connectivity and enhances sympathetic neuronal output (Brown and McAfee, 1982), defining ganglionic long-term potentiation (gLTP) as a peripheral feed-forward plasticity mechanism. It is less clear, however, how homeostatic mechanisms may constrain feed-forward processes to prevent runaway synaptic strengthening of sympathetic circuits.

The maintenance of peripheral sympathetic neuronal (SN) output is essential for the development of peripheral organs and structures (Borden et al., 2013; Kreipke and Birren, 2015) as well as their ongoing function (Kumari et al., 2024). Although sympathetic circuits such as the sympatho-cardiac pathway maintain continuous low-level activity (Barman and Yates, 2017), sustained increases in SN activity underlie diseases such as neurogenic hypertension (Pappas et al., 1974). In the spontaneously hypertensive rat (SHR), a model of neurogenic hypertension, SN overactivity precedes and drives the onset of pathological dysfunction (Magee and Schofield, 1992; Shanks et al., 2013; Davis et al., 2020; Haburčák et al., 2022). Increases in SN activity are driven by peripheral, as well as central (Liang et al., 2016), mechanisms suggesting that compensatory pathways may also be deployed in peripheral ganglia to tightly control SN output during normal development and function.

Sympathetic satellite glial cells (SGCs) regulate SN development and function (Birren et al., 2025), and are potential mediators of SN plasticity. Found in both autonomic and sensory ganglia, SGCs closely envelop neuronal somata to create a perineuronal space critical for extracellular signaling and regulation of neuronal properties (Hanani and Spray, 2020). Sympathetic SGCs also engulf synaptic inputs and regulate the development and function of synaptic sites (Hanani, 2010; Enes et al., 2020). The SGCs respond to acetylcholine (ACh) and ATP via cholinergic and purinergic receptors to induce intracellular calcium spikes that initiate downstream signaling (Kumagai and Saino, 2001; Xie et al., 2017; Feldman-Goriachnik et al., 2018).

Acute chemogenetic activation of sympathetic SGCs in mice increases SN activity and cardiac function. Conversely, prolonged SGC activation results in a decrease in blood pressure (Xie et al., 2017). Since enhanced sympathetic activity is associated with elevations in blood pressure (Seravalle and Grassi, 2022), this suggests that, over the long-term, SGCs may exert homeostatic control over SN output. Further support for this idea comes from experiments in which depletion of SGCs led to fewer SNs overall, but increased activity of the remaining neurons (Mapps et al., 2022). These studies suggest that SGCs play both a developmental and homeostatic role in the sympathetic system.

SNs have dual transmitter properties *in vivo* and *in vitro* (Furshpan et al., 1976; Landis and Keefe, 1983; Lockhart et al., 1997; Kanazawa et al., 2010), allowing for cholinergic collaterals within the sympathetic ganglia (Clyburn et al., 2023) and noradrenergic projections to the periphery (McCorry, 2007). Cultured SNs form a well-characterized cholinergic network (O’Lague et al., 1974; Gingras and Ferns, 2001; Slonimsky et al., 2003) that we used to identify a homeostatic mechanism regulating SN synaptic density in response to prolonged changes in SN and SGC activity. We determined that SGCs are essential for the remodeling of cholinergic inputs to SNs by showing that neuronal stimulation induced compensatory structural plasticity only when SGCs were present. Further, cholinergic activation of SGCs regulated glial expression of synaptic regulatory factors and was sufficient to induce compensatory changes in synaptic density. Finally, SGCs derived from SHRs show aberrant modulation of synaptic properties. This defines a new form of peripheral structural homeostatic plasticity that normally constrains sympathetic activity through compensatory changes in synaptic density.

## Materials and Methods

### Animals

All experiments using rats were approved by the Brandeis Institutional Animal Care and Use Committee. Wistar-Kyoto (WKY) and SHR rats were maintained in an in-house breeding colony. All primary cultures originated from postnatal (P0-4) rats from both male and female animals.

### Histology

The superior cervical ganglion (SG) of WKY and SHR postnatal (P0-P4) and adult rats (P150) were dissected and immediately drop-fixed in 4% paraformaldehyde (PFA, Fisher Scientific j61899-AP, Waltham, MA, USA) for one hour while rotating at 4**°**C. SGs were washed with phosphate-buffered saline (PBS) and suspended in 30% sucrose solution overnight. SGs were embedded in O.C.T compound (Fisher Scientific, 23-730-571) in a disposable mold (Fisher Scientific, 22-363-553) and stored at −80**°**C. Samples were held at −20**°**C for one hour prior to sectioning on a cryostat (Leica Biosystems CM3050s, Nussloch, Germany) set to 14µm. Sections were thaw mounted to Colorfrost plus microscope slides (Fisher Scientific 1255025) and stored at −20**°**C.

### Primary cell culture

Primary cell cultures were generated from WKY and SHR postnatal SGs as previously described (Haburčák et al., 2022). Briefly, SGs were incubated in a collagenase type I (Worthington Biomedical Corporation, LS004194, Lakewood, NJ, USA), dispase II (Gibco BRL, 17105041, Invitrogen, Carlsbad, CA, USA) enzyme mix and passed through multiple fire-polished pipettes to dissociate SGs into a single-cell suspension. Cultures used for immunocytochemistry were plated onto glass-bottomed 35mm petri dishes (Mattek P35G-1.5-14C, Ashland, MA, USA) at a density of 10,000 cells per dish. All RNA experiments were plated onto 60mm petri dishes (Fisher Scientific FB012925) at a density of 150,000 cells per dish. Culture dishes were coated with Collagen type II (50ug/ml; BD Biosciences, Bedford, MA, USA) and mouse laminin (5ug/ml; BD Biosciences) and stored at 4**°**C for 24 hours. Mouse 2.5S nerve growth factor (NGF, 5ng/ml, Alomone Labs, N-240, Jerusalem, Israel) was added to cultures to support neuronal survival and was withheld to generate isolated SGC cultures. Cytosine arabinofuranoside (AraC, Sigma C1768, 1uM, Sigma, St. Louis, MO, USA), an inhibitor of cell division, was added to cultures at 1 day in vitro (DIV) to obtain glia-free, isolated SN cultures, or at 6 DIV to prevent SGCs from overgrowing in neuron-glia co-cultures. Cell cultures were maintained by replacing half of the medium with fresh medium twice per week.

### Activity manipulations of cultured SNs and SGCs

Chronic SN activity manipulation was performed by adding 1uM of either acetylcholine hydrochloride (ACh) (Sigma A6625), or tetrodotoxin citrate (TTX) (Tocris 1069, Bristol, UK) starting at 12 DIV. Conditions receiving ACh were treated with an additional dose of ACh 24 hours after the initial treatment. A vehicle control condition was run in all experiments by treating cultures with sterile water.

Excitatory DREADDs (designer receptor exclusively activated by designer drugs) under the control of cell-specific promoters, were transduced via adeno-associated viruses (AAV, ADDGENE, Watertown, USA, Table 1). Viruses were added to the cultures at 5 DIV and DREADD expression was confirmed at 11 DIV by visualization of mCherry using an Olympus IX80 microscope (Evident Scientific, Tokyo, Japan). Chemogenetically manipulated cells were treated with either 10 µM clozapine-n-oxide dihydrochloride (CNO, HelloBio HB6149, Bristol, United Kingdom) or water (vehicle) for the final 2, 24, or 48 hours of the culture period. In some experiments, 1 µM minocycline hydrochloride (Sigma M9511) was added with or without CNO through bath application. All cultures were fixed with 4% paraformaldehyde (PFA, Fisher Scientific j61899-AP) for 15 minutes. In experiments where network activity was assessed using cFos immunofluorescence, culture media was fully replaced 30 minutes prior to PFA fixation.

**Table 1:** List of adenoassociated viruses (AAV) used in cell culture experiments. Titer expressed as vector genome per mL (vg/mL).

| **Virus** | **Supplier**<br>**(Cat.**<br>**Number)** | **Serotype** | **Titer (vg/mL)** |
| --- | --- | --- | --- |
| AAV-hSyn-hM3D(Gq)-mCherry | Addgene<br>(50474) | AAV5 | $7.5 \times 10^9$ |
| AAV-GFAP-hM3D(Gq)-mCherry | Addgene<br>(50478) | AAV5 | $2.3 \times 10^{10}$ |
| AAV-hSyn- mCherry | Addgene<br>(114472) | AAV5 | $7.5 \times 10^9$ |

### Preparation of conditioned medium

Conditioned medium (CM) was generated as previously described(Enes et al., 2020) from SN-SGC cocultures (NGCM) and isolated SGCs (GCM). Briefly, cultures were transfected with excitatory DREADDs and stimulated with CNO or vehicle for 48 hours at the end of the culture period. Media was collected after 24 and 48 hours of treatment and centrifuged at 1750xg for 90 minutes in a centrifugal concentrator (Sartorius VS15RH11, Bohemia, NY, USA) with a size cut-off filter of 5 kDA to achieve a 15x concentration. The collected media was syringe filtered and frozen. Before use, the NGCM-Veh, NGCM-CNO, GCM-Veh and GCM-CNO was diluted 1:3 in fresh medium and used to treat isolated cultures of SNs for 48 hours.

### RNA extraction and RT-qPCR

RNA was extracted from SGC and SN-SGC cultures using a DirectZol RNA micro prep kit (Zymo-research R2060, Irvine, CA, USA) and the RNA was evaluated for purity by using a nanodrop (Thermofisher Fisher Scientific, Waltham, MA, USA) and gel electrophoresis (not shown). RNA was diluted to 100ng/ul and stored at −80**°**C. RNA was converted to cDNA using an iScript cDNA synthesis kit (BioRad 1708897, Hercules, CA, USA).

The cDNA was diluted 1:30 in RNAase-free water, mixed with Sybr green PCR mix (PCR Biosystems PB20.15-05, London, England) and amplified using custom forward and reverse primers (Table 2) in an Eppendorf Mastercycler Epgradient S (Eppendorf, Hamburg, Germany). The cycle threshold was determined for the target and the reference gene GAPDH. A 2ΔΔCT2 method was used to calculate the relative gene fold change for a minimum of two technical and three biological replicates for each experiment.

**Table 2:**
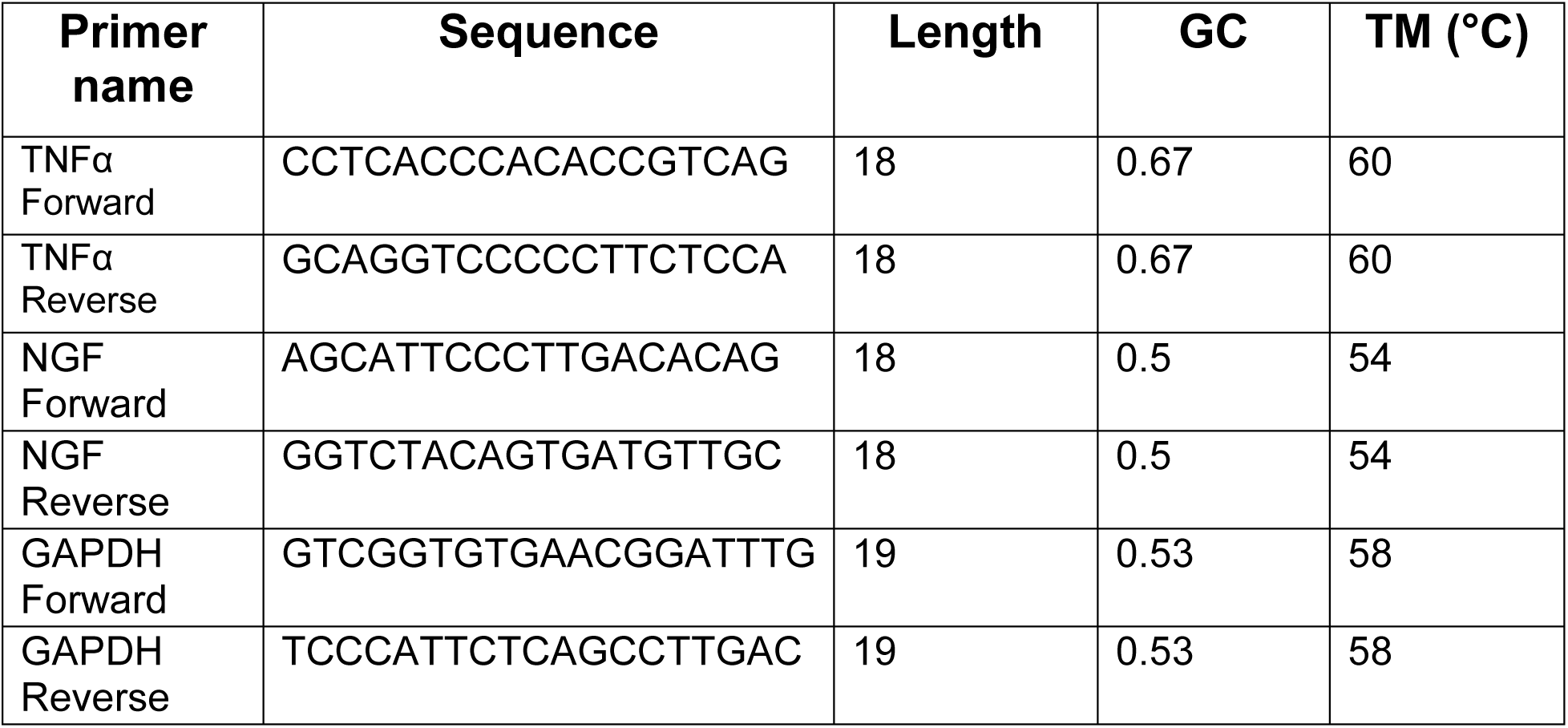
List of primers used to quantify relative RNA expression.

### Immunostaining

Cryosections of SGs were permeabilized by treatment with 1% Triton X-100 for 1 hour. Slides were washed and incubated in a blocking buffer (5% goat serum (Thermo Fisher Scientific 16260-64), 1% bovine serum albumin (BSA, Sigma A9647), 0.3% Triton-x (Sigma T8787)) for 1 hour. Samples were incubated overnight with primary antibody (Table 3) at room temperature. Slides were washed in PBS and incubated overnight in secondary antibody (Table 3) at room temperature. Samples were washed and mounted using Prolong glass antifade mountant (Invitrogen 3112801, Waltham, MA, USA).

**Table 3:** List of antibodies used for immunostaining.

| Primary antibody | Supplier (Cat. Number) | RRID | Host | Dilution |
| --- | --- | --- | --- | --- |
| VACHT | Synaptic Systems (139103) | AB_887864 | Rabbit | 1:1000 |
| Shank | Millipore (MABN24) | AB_11213314 | Mouse | 1:500 |
| MAP2 | Millipore (AB5543) | AB_571049 | Chicken | 1:1000 |
| s100β | Invitrogen (MA1-25005) | AB_795382 | Mouse | 1:500 |
| RFP | Invitrogen (PA1-986) | AB_560939 | Rabbit | 1:1000 |
| Tyrosine Hydroxylase | Abcam (AB76442) | AB_1524535 | Chicken | 1:1000 |
| nAChR3 | DSHB (6A12-11c) | AB_2721952 | Mouse | 1:500 |
| cFos | Abcam (222699) | AB_2891049 | Rabbit | 1:250 |

| Secondary | Supplier (Cat. Number) | RRID | Host | Dilution |
| --- | --- | --- | --- | --- |
| Alexa fluor 488 | Invitrogen (A11029) | AB_2534088 | Goat | 1:1000 |
| Alexa fluor 568 | Invitrogen (A32733) | AB_2866492 | Goat | 1:1000 |
| Alexa fluor 647 | Invitrogen (A11031) | AB_144696 | Goat | 1:1000 |

Fixed cultures were permeabilized with 1% Triton-x for 10 minutes and washed in blocking buffer containing 5% Goat serum, 1% BSA, 0.025% Triton-X. Cultures were incubated in primary antibody solution (Table 3) overnight at 4**°**C. After washing, dishes were incubated with secondary antibody (Table 3) overnight at 4**°**C. For cFos analysis, cultures were co-stained with MAP2 (Table 3) and treated with 0.1ug/ml DAPI (Sigma D9542) for 5 minutes after secondary antibody treatment.

### Imaging and image analysis

All images were captured on a Zeiss LSM 880 confocal microscope (Oberkochen, Germany) with a 63X objective. Microscope settings were kept consistent across imaging sessions, with all images captured as 8-bit and 1024×1024 in size. Confocal images were taken using a z-stack setting that captured images at 0.7 µm steps with no changes to laser channel settings during imaging sessions. For culture experiments, coverslips were divided into quadrants and a minimum of 15 SNs were captured for each condition across all four quadrants. The MAP2 channel was used to find and center the image on the cell body of individual SNs and to determine the z-stack range. In histology experiments, as well as cultures stained for cFos, images were captured by tiling across the sample. A maximum intensity projection was generated for image analysis using Zeiss Zen Microscopy Software (RRID:SCR_013672, Oberkochen, Germany).

Synaptic quantification of maximum intensity projections of cultures and cryostat sections was performed in Puncta Analyzer, an ImageJ plugin (Ippolito and Eroglu, 2010) as described (Haburcak et al., 2022). This software calculates the center of mass for each puncta and considers a colocalized marker to be any pre- and post-synaptic puncta that is overlapped within a pre-defined pixel distance. Colocalized and synaptic marker counts were normalized to the MAP2 area. In culture experiments, a minimum of 3 independent cell culture preparations were analyzed. In histological experiments, a minimum of three cryosections from the right SG of at least 6 male animals per strain were analyzed and averaged. The relative glial effect on synaptic density in WKY and SHR cultures was determined by subtracting the averaged synaptic density of isolated SNs (N_av_) from the synaptic density in SN+SGC (NG) cocultures and dividing by the N_av_ density for each independent cell culture preparation ((NG-N_av_)/N_av_). Cellprofiler (version 4.2.6, Broad Institute, Cambridge, Massachusetts, USA) software was used to quantify the corrected total cell fluorescence of cultured SGCs stained for s100β. For cFos analysis, conditions were blinded and manually counted using Zen Blue image analysis software by calculating the percentage of cFos positive nuclei in MAP2-positive SNs and comparing to DAPI-positive nuclei.

### Statistics

All immunostaining data involving multiple comparison were analyzed using a 1-way (for a single variable) or 2-way (for two variables) ANOVA with a post-hoc Sidaks or Tukey’s test unless otherwise stated. An unpaired t-test was used for comparing colocalized synaptic density between two conditions and a paired t-test was used for analysis of relative RNA expression.

All experimental data was plotted using Prism software (GraphPad; San Diego, CA, USA) and figures were assembled in Adobe Illustrator (Adobe, Inc; San Jose, CA, USA). The diagram was prepared using BioRender (BioRender.com, Ontario, Canada). All results are presented as mean ± S.E.M. Statistical analysis was performed using Prism. P values are represented as *<0.05. **<0.01. ***0.001, ****0.0001, or n.s. (non-significant).

## Results

### Homeostatic regulation of sympathetic neuron synaptic density following prolonged activity manipulation

We investigated the effects of prolonged activity manipulations on cholinergic synaptic density in sympathetic ganglionic cultures containing SNs and SGCs. Cultures were grown for 14 days and chronically treated with ACh for the last 48 hours of the culture period (Fig. 1A). Cultures were fixed and stained for the presynaptic marker Vesicular Acetylcholine Transporter (VAChT), the postsynaptic marker Shank and the neuronal cell body and dendritic marker Microtubule-associated protein 2 (MAP2) (Fig. 1B). Synaptic density was measured as the number of colocalized pre- and postsynaptic sites per area of MAP2+ staining. ACh-stimulated SNs had a decreased density of colocalized synaptic markers compared to vehicle treated cultures (Fig. 1C left). This was associated with a downregulation in the number of VAChT, but not Shank puncta (Fig. 1C right), suggesting a presynaptic contribution to activity-dependent regulation of synaptic sites.

**Figure 1:**
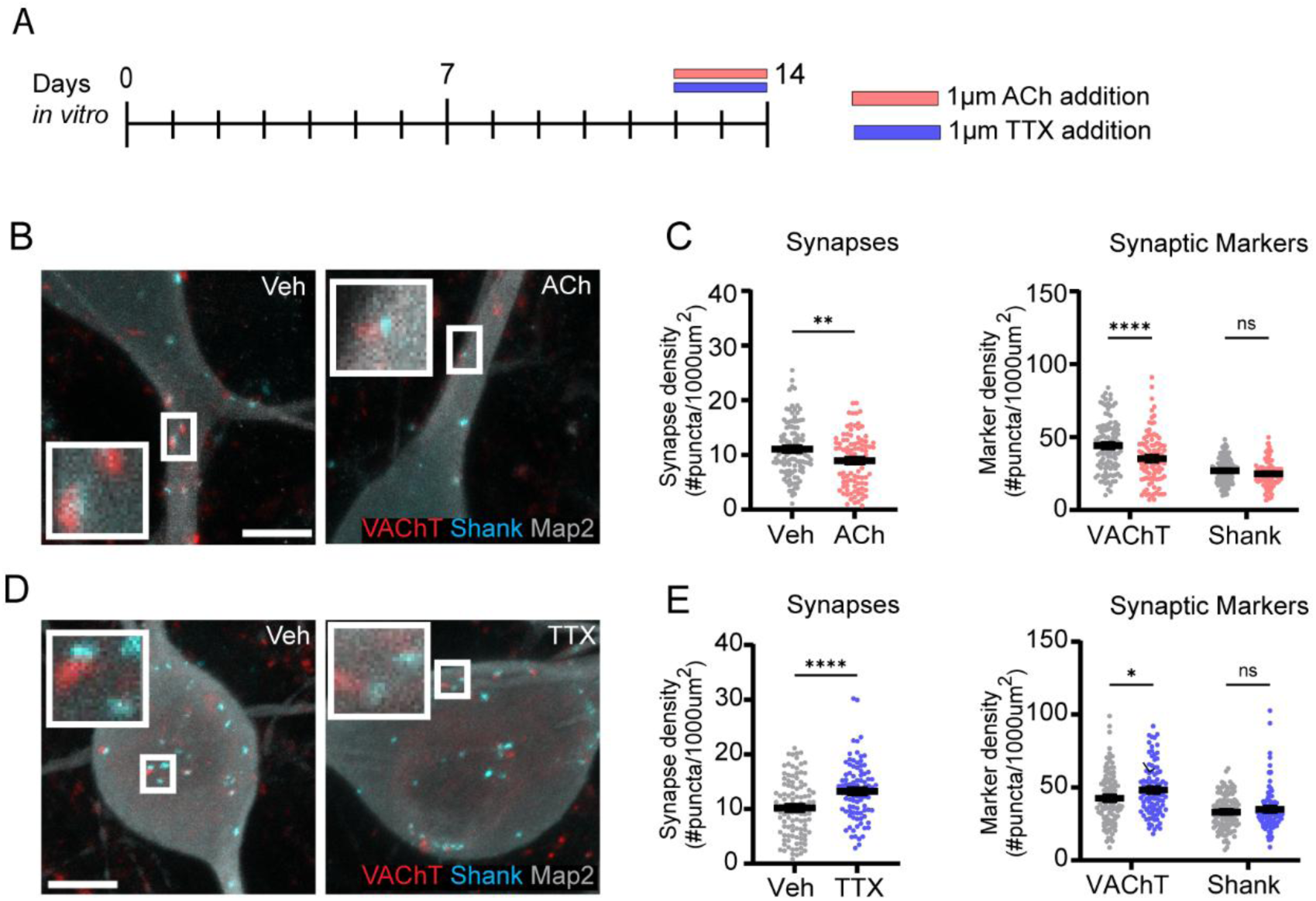
Chronic activity manipulation of SNs induces homeostatic changes in synaptic structural properties. **A,** Experimental setup for pharmacological treatment of SNs in ganglionic cultures with TTX or ACh on DIV 12 and 13. Legend colors are consistent across all analyses. **B,** Representative images of SNs treated with Vehicle (Veh) or 1 µM ACh and stained for VAChT, Shank, and MAP2 (Scale = 10 µm). Examples of colocalized synaptic markers (large white box) taken from a magnified region of interest (Small white box). **C,** Quantification of synaptic density (left) and individual pre- and postsynaptic markers (right) following ACh treatment (n=number of neurons; Veh n=108, ACh n=96. **Left**: **p=0.0028. **Right**: VAChT ****p<0.0001, Shank n.s.). **D,** Representative images of SNs treated with Veh or 1 µM TTX stained for VAChT, Shank, and MAP2 (Scale = 10 µm). **E,** Quantification of synaptic density (left) and pre- and postsynaptic markers (right) following TTX treatment. (Veh n=106, TTX n=95; **Left**: TTX **** p<0.0001, **Right**: VAChT *p=0.0228, Shank n.s.).

To determine if the compensatory decrease in synaptic density was part of an activity-dependent homeostatic regulatory system, we next inhibited neuronal activity with the voltage-gated sodium channel blocker tetrodotoxin (TTX) for 48 hours (Fig. 1D). Cocultured SNs chronically inhibited with TTX had a higher synaptic density than vehicle-treated cultures (Fig. 1E left). Presynaptic, but not postsynaptic, marker expression was also upregulated (Fig. 1E right) in TTX-treated neurons, further suggesting a presynaptic contribution to homeostatic shifts in synaptic density.

We further investigated activity-dependent regulation of synaptic sites by introducing an excitatory Designer Receptor Exclusively Activated by Designer Drugs (DREADDs) selectively into the cocultured SNs using an AAV-Syn-hM3D(Gq) virus (Fig. 2A-B). Cultures were treated with clozapine-n-oxide (CNO) or water (vehicle) for the last 2, 24, or 48 hours of the 14-day culture period and stained for colocalized synaptic sites (Fig. 2A, C). Synaptic density decreased following CNO treatment at 24- and 48-hours, with no change in the 2-hour condition (Fig. 2D left). Similar to SN activation with ACh, presynaptic, but not postsynaptic puncta were downregulated after 24- and 48-hours of CNO exposure. In contrast to the longer time-points, presynaptic marker expression was significantly upregulated at the 2-hour time point (Fig. 2D right), suggesting a potential early effect on synaptic properties.

**Figure 2:**
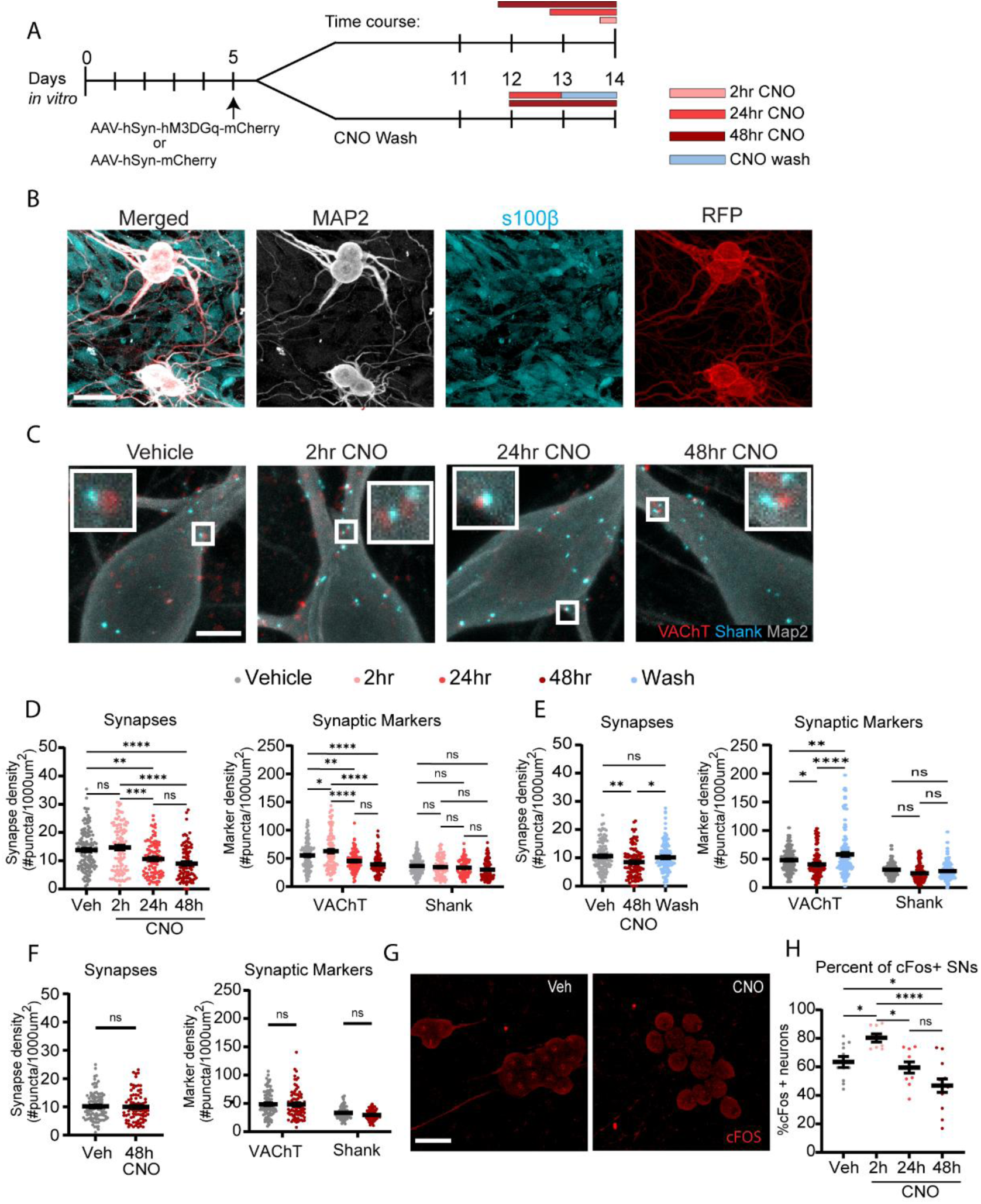
Chemogenetic manipulation of SNs induces downregulation of SN synaptic structural properties and activity. **A,** Experimental setup for chemogenetic activation of SNs in ganglionic cultures by treatment with either vehicle (Veh) or 10 µM CNO for 2, 24, or 48 hours followed by staining for synaptic markers or cFos (top) or for a CNO wash experiment (bottom). **B,** Representative images of postnatal WKY SNs treated with Veh or CNO and stained for MAP2, s100β, or red fluorescent protein (RFP) (Scale = 10 µm). **C,** Representative images of WKY SNs treated with Veh or CNO and stained for VAChT, Shank, and MAP2 (Scale = 10 µm). **D,** Quantification of synaptic density (left) and individual synaptic markers (right) in chronically stimulated SNs in ganglionic cultures. (n=number of neurons, Veh n=135, 2hr n=105, 24hr n=97, 48hr n=92. **Left**: Veh vs. 2hr n.s; Veh vs. 24h **p=0.0041; Veh vs. 48hr **** p<0.0001; 2hr vs. 24h *** p=0.0002; 2hr vs. 48hr ****p<0.0001; 24h vs. 48hr n.s. **Right**: Veh vs. 2hr VAChT *p=0.0211, Shank n.s; Veh vs. 24hr VAChT **p=0.0019, Shank n.s.; Veh vs. 48hr VAChT **** p<0.0001, Shank n.s.; 2hr vs. 24hr VAChT **** p<0.0001, Shank n.s.; 2hr vs. 48hr VAChT **** p <0.0001, Shank n.s.; 24hr vs. 48hr VAChT n.s., Shank n.s.). **E,** Quantification of synaptic density (left) and individual synaptic markers (right) for SNs following a 24 hour wash. (Veh. n=112, CNO n=105, Wash n=110. **Left:** Veh vs 48h CNO ** p=0.0087; Veh. vs Wash n.s.; 48h CNO vs. Wash *p=0.0445. **Right:** Veh. vs 48h CNO VAChT * p=0.0403, Shank n.s; Veh. vs. Wash **p=0.0018, Shank n.s.; 48h CNO vs Wash VAChT **** p<0.0001, Shank n.s.). **F**), Quantification of synaptic density (left) and individual synaptic markers (right) for SNs with viral-vector control (no DREADD) treated with Veh or CNO. (Veh n=160, CNO n=122. **Left**: Veh vs 48h CNO n.s. **Right**: Veh vs 48hr CNO VAChT n.s., Shank n.s.). **G,** Representative images of SNs strained for cFos (red) after treatment with CNO or Veh. For 48 hours. (Scale = 50 µm). **H,** Quantification of the percentage of cFos positive SN nuclei in the cholinergic SN network after chronic activity manipulation. (n=number of culture plates: Veh. n=11, 2hr n=8, 24hr n=9, 48hr n=13. Veh. vs. 2hr * p= 0.049; Vehicle vs. 24hr n.s.; Veh. vs. 48hr * p=0.0236; 2hr vs. 24hr * p=0.0127; 2hr vs. 48hr *** p<0.0001; 24hr vs. 48hr n.s.).

We examined the reversibility of the activity-dependent process by removing CNO from the cocultures 24 hours after the initiation of treatment (Fig. 2A), a timepoint at which we observed a significant reduction in synaptic density. We found a rebound in synaptic density following the CNO washout (Fig. 2E left). While the density of colocalized sites returned to baseline following a 24-hour wash, the presynaptic marker in the washout condition was significantly higher than the baseline condition (Fig. 2E right), suggesting a compensatory overshoot of presynaptic marker expression.

We asked if there were off-target effects of CNO in our culture system by introducing a control viral vector that lacks DREADDs. CNO treatment for 48 hours did not induce a change in either synaptic density (Fig. 2F left) or individual synaptic marker number (Fig. 2F right). These findings show that the regulation of synaptic density is through a CNO-driven activity-dependent process and not through other, indirect actions of CNO.

We next examined neuronal cFos expression to ask if chronic SN stimulation influenced the ongoing activity of SNs in the cholinergic circuit. DREADD-expressing SN cocultures were treated with vehicle or CNO for 2, 24 or 48 hours and stained for cFos (Fig. 2A, G). Treatment with CNO for 2 hours resulted in an upregulation of cFos-positive neuronal nuclei (Fig. 2H) consistent with an acute effect of chemogenetic activation of the circuit. There was a significant decrease in cFos-positive SNs at 24-and 48-hours, supporting the idea that activity-dependent downregulation of synapses acts as a compensatory mechanism to normalize activity in the system.

### Satellite glial cells regulate SN circuit activity by altering synaptic structural properties

We asked if SGCs in the sympathetic cultures contributed to activity-dependent synaptic plasticity. We depleted SGCs from the cultures and introduced the excitatory DREADD into the isolated SNs and measured synaptic density following 48hrs of CNO treatment (Fig. 3A-B). In contrast to cultures that are grown in the presence of SGCs, chronically stimulated isolated SNs showed a modest but significant increase in synaptic density (Fig. 3C left) and presynaptic puncta (Fig. 3C right). This shows that in the absence of SGCs compensatory responses to activity are lost, and feed-forward changes in synaptic density are observed.

**Figure 3:**
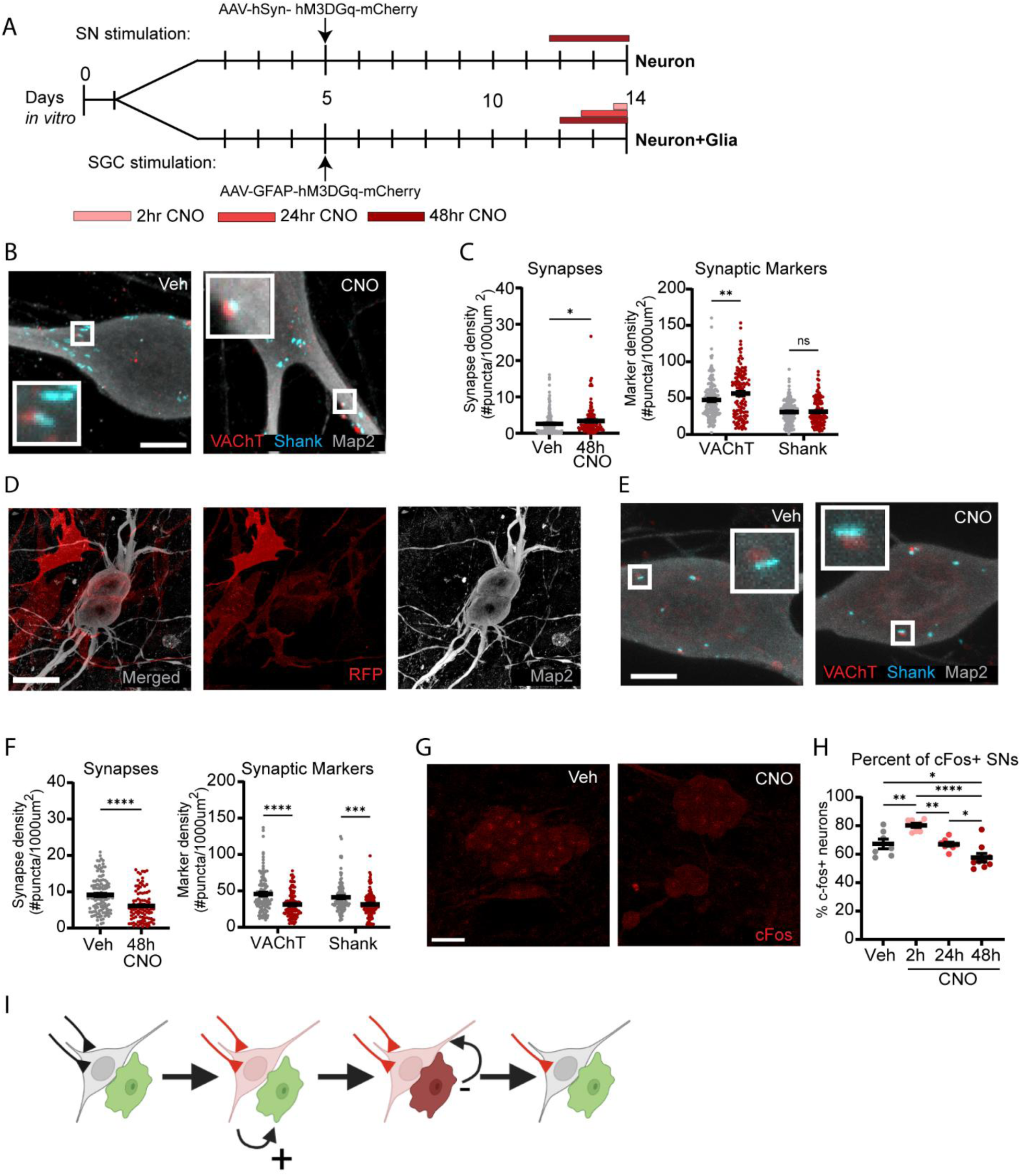
Activity-dependent synaptic compensation requires chronic glial activation. **A,** Experimental setup for chemogenetic activation of isolated SNs in the absence of glia (top) or of SGCs in ganglionic cultures (bottom). Cultures were treatment with either vehicle (Veh) or 10 µM CNO and stained for synaptic markers or cFos at the end of the culture period. **B,** Representative images of isolated WKY SNs treated with Veh or CNO and stained for VAChT, Shank, and MAP2 (Scale = 10 µm). **C,** Quantification of synaptic density (left) and individual synaptic markers (right) in chronically stimulated SNs in isolated cultures. (n=number of neurons; Veh n=160, CNO n=122. **Left**: Veh vs. 48h CNO **p=0.0439. **Right**: Veh vs. 48h CNO VAChT ** p=0.0037, Shank n.s.). **D,** Representative images of SNs with DREADDs-expressing SGCs stained for MAP2 and RFP to visualize mCherry (Scale = 10 µm). **E,** Representative images of SNs cocultured with DREADD-expressing SGCs treated with Veh or CNO for 48 hours and stained for VAChT, Shank, and MAP2 (Scale = 10 µm). **F,** Quantification of synaptic density (left) and individual synaptic markers (right) in SNs cultured with DREADD-expressing SGCs after treatment with Veh or CNO for 48 hours. (Veh n=117, CNO n=97. **Left**: Veh vs. 48h CNO ****p<0.0001. **Right**: Veh vs. 48h CNO VAChT ****p<0.0001, Shank ** p=0.0022). **G,** Representative images of SNs cultured with DREADD-expressing SGCs treated with 48 hours of Veh or CNO and stained for cFos (red). **H,** Quantification of the percentage of cFos-positive SNs in the cholinergic SN network after chronic activity manipulation of SGCs. (n=number of culture plates; Veh. n=7, 2h CNO n=8, 24h CNO n=8, 48h CNO n=9. Veh vs. 2h CNO **p= 0.048; Veh vs. 24h CNO n.s.; Veh vs. 48h CNO *p=0.0399; 2h CNO vs. 24h CNO **p=0.0029; 2h CNO vs. 48h CNO ****p<0.0001; 24h CNO vs. 48h CNO *p=0.0372). **I,** Proposed model of homeostatic regulation of SN synaptic density. SNs at baseline activity (grey) and SGCs (green) develop in close contact with cholinergic synapses (black). Chronic alterations in neuronal activation (red) activate SGCs (dark red) reducing SN synaptic structural and functional properties move the SN closer to baseline activity.

We examined the impact of glial activation on synaptic regulation of cocultured SNs by using a glial fibrillary acidic protein (GFAP) promoter to selectively express an excitatory DREADD in SGCs (Fig. 3A, D). Cultures were treated with CNO or vehicle for the last 48 hours of the 14-day culture period and stained for synaptic and neuronal markers (Fig. 3E). Chronic glial activation resulted in a significant decrease in synaptic density (Fig. 3F left), as well as a reduction in both VAChT and Shank puncta density (Fig. 3F right).

We next used cFos to examine the functional impact of SGC stimulation on the activity of the SN network. Cocultures containing DREADD-expressing SGCs were treated with vehicle or CNO for 2, 24 or 48 hours and stained for cFos expression (Fig. 3G). Two hours of CNO-treatment increased the number of cFos-positive SNs (Fig. 3H), confirming, as previously reported in mice (Xie et al., 2017), that short-term chemogenetic activation of SGCs enhances SN activity. We observed a significant decrease in cFos-positive SNs in both the 24- and 48-hour conditions compared to the 2-hour treatment, with the 48-hour treatment showing a decrease compared to the baseline vehicle control. Since SGCs are activated by cholinergic signaling (Feldman-Goriachnik et al., 2018), these data suggest a model in which SGCs act downstream of neuronal activity to modify synaptic density and function in a compensatory manner (Fig. 3I).

We used the glial inhibitor minocycline (Tikka et al., 2001) to determine if glial activation is necessary for the induction of activity-dependent plasticity (Fig. 4A-B). Minocycline treatment alone did not influence synaptic density in the absence of chemogenic excitation of SNs (Fig. 4C Left). Concurrent treatment with minocycline and CNO prevented activity-dependent downregulation of synaptic density compared to SNs treated with CNO alone (Fig. 4C left). This indicates that SGC signaling is necessary for induction of activity-dependent plasticity. While minocycline blocked activity-dependent decreases in synaptic sites, there was an overall decrease in presynaptic markers (Fig. 4C Right), suggesting a partial effect of the blocker.

**Figure 4:**
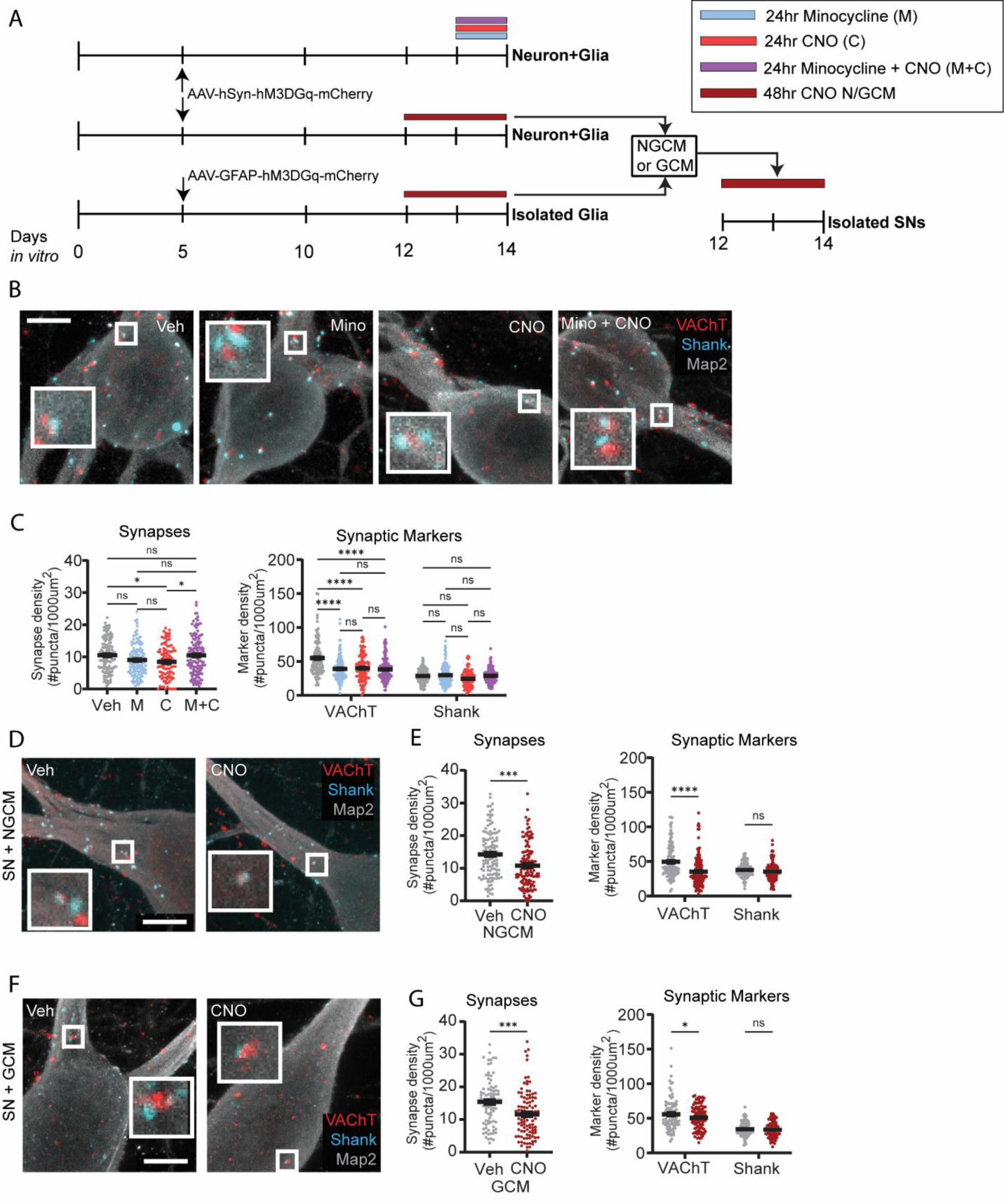
SGC activation and release of glial-derived synapse-regulatory factors induce a compensatory decrease in SN synaptic density. **A,** Experimental setup for analysis of glial-dependent regulation of synaptic density. Chemogenetically-activated SNs in coculture were treated with vehicle (Veh), 10 µM CNO, 1 µM Minocycline (Mino), or CNO and Mino (M+C) for 24 hours (top). Isolated, cultured SNs were treated with conditioned medium (CM) from chemogenetically-activated SNs in coculture (NGCM) (middle) or with CM from chemogenetically-activated isolated glia (GCM) (bottom). **B,** Representative images of SNs in cocultures treated with Veh, CNO, Mino, or M+C for 24 hours and stained for VAChT, Shank, and MAP2 (Scale = 10 µm). **C,** Quantification of synaptic density (left) and individual synaptic markers (right) in SNs in cocultures treated with Veh, CNO, Mino, or M+C for 48 hours. (n=number of neurons; Veh n=160, CNO n=122. **Left**: Veh vs. CNO *p=0.0232; Veh vs. Mino n.s.; Veh vs. M+C n.s.; CNO vs. Mino n.s.; CNO vs. M+C * p=0.0211; Mino vs. M+C n.s. **Right:** Veh. vs. CNO VAChT**** p<0.0001, Shank n.s.; Veh. vs. Mino VAChT **** p<0.0001, Shank n.s.; Vehicle vs. M+C VAChT **** p<0.0001, Shank n.s.; CNO vs. Mino VAChT n.s, Shank n.s.; CNO vs. M+C VAChT n.s., Shank n.s.; Mino vs. M+C VAChT n.s., Shank n.s.). **D,** Representative images of isolated, cultured SNs treated with NGCM-Veh or NGCM-CNO for 48 hours and stained for VAChT, Shank, and MAP2 (Scale = 10 µm). **E,** Quantification of synaptic density (left) and individual synaptic markers (right) in isolated, cultured SNs treated with NGCM-Veh or NGCM-CNO for 48 hours. (NGCM-Veh n=120, NGCM-CNO n=129. **Left**: NGCM-Veh vs. NGCM-CNO ***p<0.0001. **Right**: NGCM-Veh vs. NGCM-CNO VAChT ****p<0.0001, Shank n.s.). **F,** Representative images of isolated, cultured SNs treated with GCM-Veh or GCM-CNO for 48 hours and stained for VAChT, Shank, and MAP2 (Scale = 10 µm). **G,** Quantification of synaptic density (left) and individual synaptic markers (right) in isolated SNs treated with GCM-Veh or GCM-CNO for 48 hours. (GCM-Veh n=89, GCM-CNO n=101. **Left**: GCM-Veh vs GCM-CNO ***p=0.0004. **Right**: GCM-Veh vs GCM-CNO VAChT *p=0.0421, Shank n.s.).

We next investigated if SGCs regulate SN synaptic density through the release of soluble glial factors. We generated concentrated conditioned medium from neuron-glial cocultures (NGCM) in which the DREADD was selectively expressed in the SNs and treated with vehicle (NGCM-Veh) or CNO (NGCM-CNO) for the final 48 hours of the culture period (Fig. 4A, D). This NGCM (CNO or vehicle-treated) was used to treat SNs grown in the absence of SGCs for 48 hours at the end of the 14-day culture. Cultures treated with NGCM-CNO showed a reduction in synaptic density compared to NGCM-Veh (Fig. 4E Left). We also observed a decrease in presynaptic, but not postsynaptic marker expression (Fig. 4E Right), showing that activated SGCs release a regulatory factor(s) that influences synaptic density and the expression of cholinergic presynaptic proteins.

The presence of activated neurons in the cocultures used to generate NGCM raises the question of whether synaptic regulatory factors were derived from the SNs or SGCs. We therefore asked if treatment of SN-only cultures with conditioned medium from chemogenetically activated isolated SGCs (GCM) also reduces synaptic sites (Fig. 4A). Isolated SNs were treated with GCM-CNO or GCM-Veh for 48 hours and stained for cholinergic synaptic markers (Fig. 4F). SNs that received GCM-CNO had decreased synaptic density (Fig. 4G left) and presynaptic VAChT expression (Fig. 4G right) compared to SNs treated with GCM-Veh. These data suggest that chronic activation of SGCs triggers regulation of glial factors that regulate synaptic machinery.

Nerve growth factor (NGF) regulates sympathetic synaptic structure and function (Lockhart et al., 1997; Tsui-Pierchala and Ginty, 1999; Lockhart et al., 2000; Luther and Birren, 2009; Luther et al., 2013; Enes et al., 2020) and is released by SGCs, as well as by targets of sympathetic innervation (Levi-Montalcini, 1964). We asked if expression of NGF was modulated in an activity-dependent manner in cocultures of SNs and SGCs. The excitatory DREADD was selectively expressed in the SNs, and cultures were treated with CNO or vehicle for 48 hours followed by quantification of NGF RNA using RT-qPCR (Fig. 5A-B). CNO-treated cocultures had reduced expression of NGF RNA (Fig. 5B) compared to vehicle-treated cultures, suggesting that glial NGF production is inversely related to cholinergic SN activity, consistent with the reduction in synaptic density seen in CNO-cultures.

**Figure 5:**
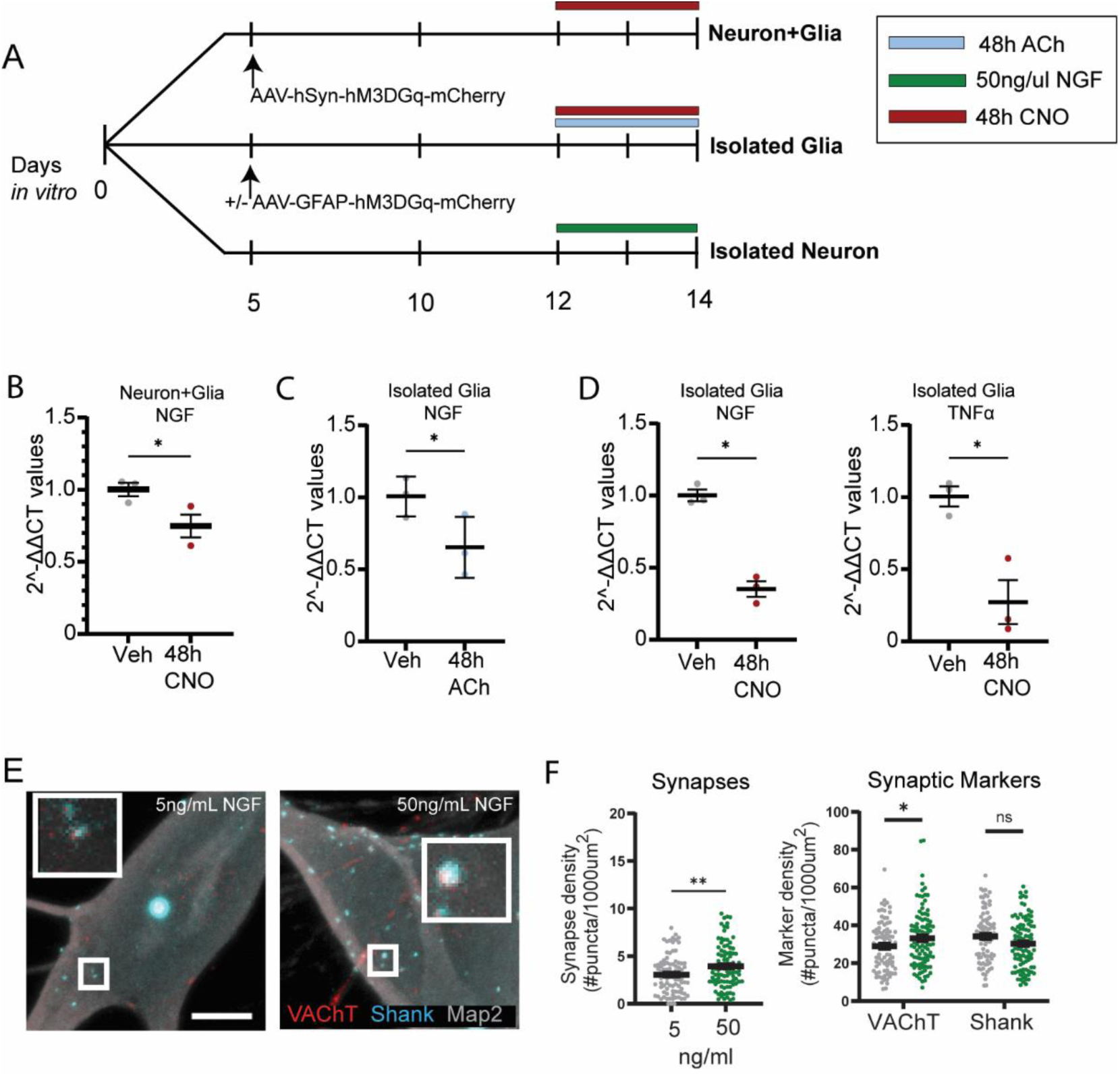
Synaptic regulatory genes are downregulated in SGCs following chronic chemogenetic or cholinergic stimulation. **A,** Experimental setup for measuring relative RNA expression and synaptic density in culture. DREADD-expressing SNs cocultured with SGCs were treated with vehicle (Veh) or CNO for 48 hours (top). Isolated, cultured SGCs were either chemogenetically stimulated by treatment with Veh or CNO, or were pharmacological stimulated with 50 µM ACh, for 48 hours (middle). Isolated, cultured SNs were treated with 5ng/mL (control) or 50ng/mL NGF for 48 hours (bottom). **B,** Relative NGF RNA expression in DREADD-expressing SNs in cocultures treated with Veh or CNO for 48 hours. (n=Independent cultures; Veh n=3, 48g CNO n=3. Veh vs. 48h CNO *p=0.0490). **C,** Relative NGF RNA expression in isolated SGCs treated with Veh or ACh for 48 hours. (Veh n=3, 48h ACh n=3. Veh vs. 48h ACh NGF: *p=0.0200). **D,** Relative NGF (Left) and TNFα (Right) RNA expression in DREADD-expressing SGCs treated with Veh or CNO for 48 hours. (Veh n=3, CNO n=3. **Left:** Veh vs. 48h CNO NGF *p=0.0202). **Right:** Veh vs. 48h CNO TNFα *p=0.0414). **E,** Representative images of isolated, cultured SNs treated with 5 ng/mL or 50 ng/mL NGF for 48 hours and stained for VAChT, Shank, and MAP2 (Scale = 10 µm). **F,** Quantification of synaptic density (left) and individual synaptic markers (right) in isolated SNs treated with 5 or 50 ng/mL NGF for 48 hours (n=number of neurons; 5 ng/ml NGF n=89, 50 ng/ml NGF n=98. **Left**: 5 ng/mL vs. 50 ng/mL **p=0.0047. **Right**: 5 ng/mL vs. 50 ng/mL VAChT *p=0.0376, Shank n.s.).

To confirm that NGF was specifically regulated in SGCs by cholinergic signaling, we treated isolated SGCs for 48 hours with ACh or vehicle (Fig. 5A). ACh-treated SGCs had reduced expression of NGF RNA (Fig. 5C). We asked if this regulation took place downstream of SGC activation by expressing excitatory DREADDs in isolated SGCs (Fig. 5A). NGF RNA expression was reduced in CNO-treated SGCs compared to vehicle-treated cultures (Fig. 5D left). Together, these data suggest a model in which cholinergic transmission in a sympathetic circuit reduces the levels of a synapse promoting factor(s) in neighboring SGCs, contributing to compensatory activity-dependent plasticity in SNs.

Tumor necrosis factor-α (TNFα) is a proinflammatory cytokine that has been implicated in glial-induced neuronal plasticity mechanisms in the CNS (Heir et al., 2024). Peripherally, TNFα is expressed in the sympathetic and sensory systems, where it promotes increased neuronal activity (Song et al., 2014; Wu et al., 2025). We examined the expression of TNFα RNA in isolated SGC cultures treated with CNO or vehicle for the final 48 hours of the culture period. CNO-stimulated SGCs showed decreased TNFα RNA compared to the vehicle control (Fig. 5D right), suggesting that SGCs may release multiple regulatory molecules that work either individually or coordinately to mediate plasticity in this system.

We reasoned that if a reduction in NGF contributed to downregulation of synaptic density, increasing NGF levels would increase synapse number. We treated isolated SN cultures with 50ng/mL of NGF for 48 hours, a concentration known to influence sympathetic neurotransmitter properties (Lockhart et al., 1997), and measured colocalized synaptic sites (Fig. 5E). SNs that received the higher concentration of NGF showed an increase in synaptic density (Fig. 5F left) and presynaptic marker expression (Fig. 5F right) compared to control SNs, supporting the idea that modulation of NGF release by SGCs provides a potential mechanism for structural synaptic plasticity.

These data collectively suggest a model where SGCs contribute to the development and maintenance of cholinergic connections between SNs (Figs 3,4, see also (Enes et al., 2020)). The SGCs respond to cholinergic signaling via muscarinic receptors (Hanani, 2010; Feldman-Goriachnik et al., 2018), allowing ongoing monitoring of synaptic transmission within the circuit. In this model, a chronic increase in neuronal activity and glial activation leads to the downregulation of synapse-promoting factors such as NGF and TNFα (Fig. 6B). Decreased availability of these released factors triggers a decrease in cholinergic synaptic density resulting in homeostatic stabilization of SN circuit activity (Fig. 6C).

**Figure 6:**
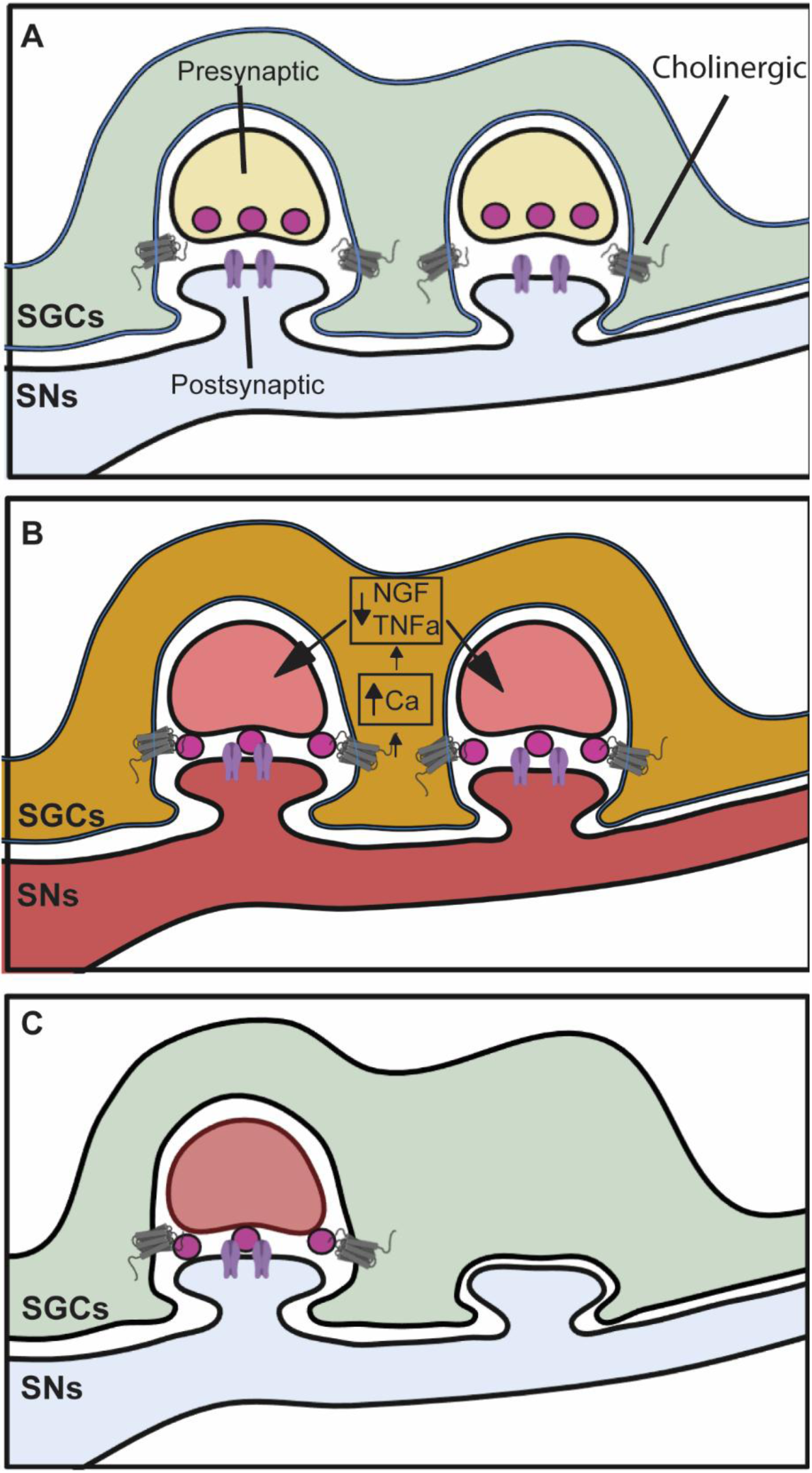
Model for activity-dependent synaptic compensation in SNs. **A,** SGCs (green) develop tight connections with SNs synaptic inputs (yellow/blue) and can be activated by cholinergic neuronal transmission. **B,** Persistent stimulation of SNs (red) increases intracellular SGC calcium leading to downregulation of soluble release factors such as TNFα and NGF. **C,** A decrease in SGC (green) release of soluble factors initiates downregulation of presynaptic cholinergic sites (blue), resulting in decreased synaptic density and SN activity.

### Satellite glial signaling and cholinergic synaptic development is disrupted in the spontaneously hypertensive rat

We next examined SN and SGC interactions in the SHR strain, an *in vivo* model of chronically increased sympathetic activity in which heightened sympathetic drive results in the development of hypertension (Brock et al., 1996; Dickhout and Lee, 1998; Shanks et al., 2013; Haburčák et al., 2022). We quantified cholinergic synapses in cryosections of the superior cervical ganglia of pre-hypertensive SHR and control WKY rats (Fig. 7). Cholinergic synapses were visualized in P2-4 ganglia by labeling for presynaptic VAChT, postsynaptic nicotinic acetylcholine receptor subunit 3 (nAChR3), and MAP2 (Fig. 7A). The density of colocalized pre-and postsynaptic sites associated with MAP2-labelled SNs, as well as the total density of the individual markers, was quantified and compared between the two strains. Postnatal SHR ganglia had a lower synaptic density than the WKY (Fig. 7B left), with non-significant changes in the number of pre- and post-synaptic markers (Fig. 7B right).

**Figure 7:**
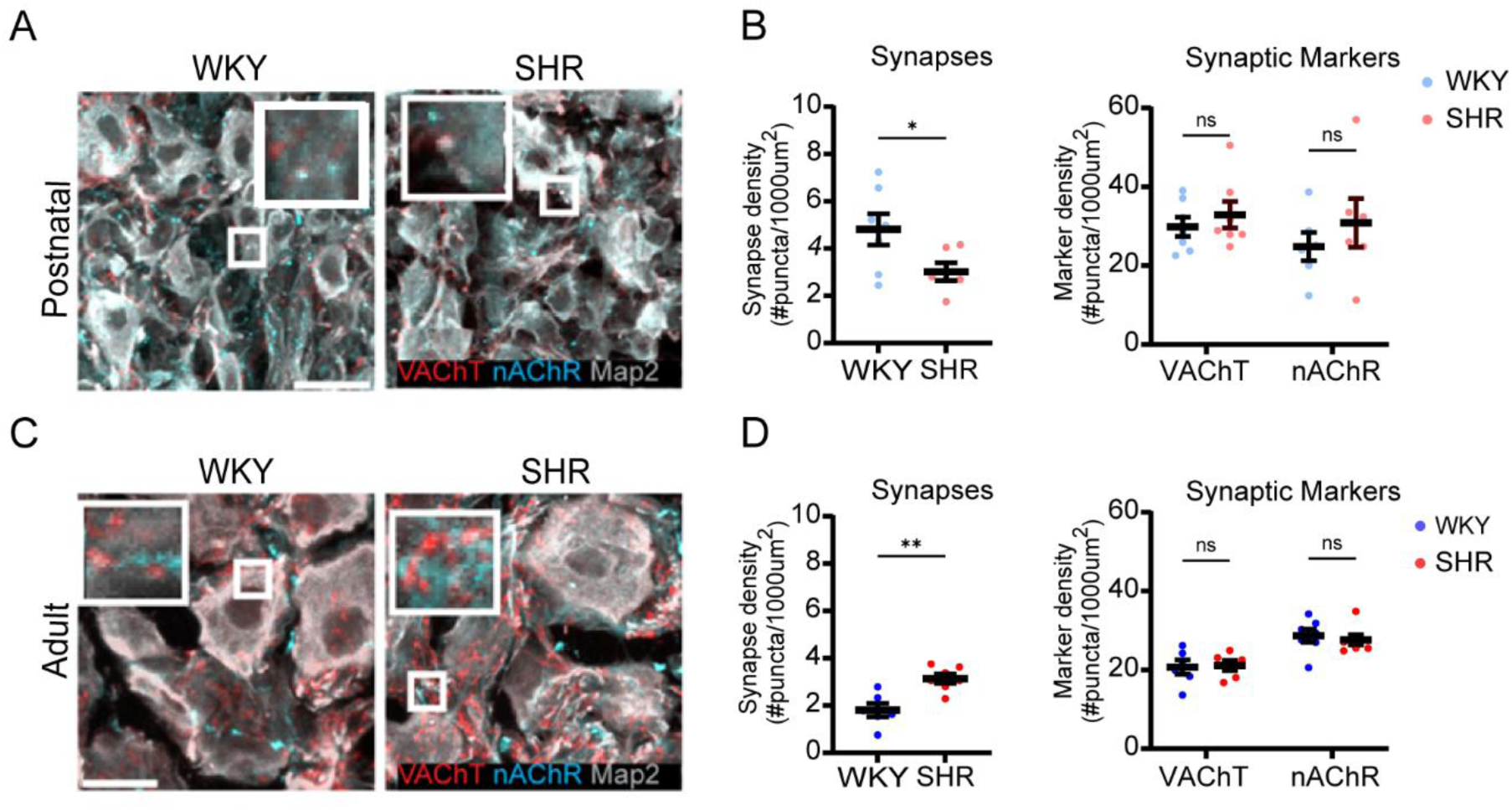
Developmental changes in synaptic density in pre-hypertensive and hypertensive SHRs. **A,** Representative images of postnatal WKY (left) and SHR (right) ganglionic cryosections stained for VAChT, nAChR α3, and MAP2. (Scale = 10 µm). **B,** Quantification of synaptic density (left) and individual pre- and postsynaptic markers (right) of WKY and SHR ganglionic sections. (n=number of animals; WKY n=7, SHR n=6. **Left**: WKY vs SHR *p=0.0474. **Right**: WKY vs. SHR VAChT n.s., Shank n.s.). **C,** Representative images of sections from adult WKY (left) and SHR (right) ganglia stained for VAChT, nAChR3, and MAP2. (Scale = 10 µm). **D,** Quantification of synaptic density (left) and individual pre- and postsynaptic markers of adult ganglionic sections. (WKY n= 6, SHR n=7. L**eft**: WKY vs SHR **p=0.0022. **Right**: WKY vs. SHR VAChT n.s., Shank n.s).

We asked if the decrease in cholinergic synapses in the young SHR ganglia was lost as the animals aged and became hypertensive. We examined cholinergic synaptic density in ganglionic cryosections from hypertensive adult (P150) SHR and normotensive WKY rats (Fig. 7C). In contrast to the young animals, adult SHRs have significantly higher synaptic density than the normotensive controls (Fig. 7D left) with no change in individual synaptic markers between the WKY and SHR (Fig. 7D right). Together, these data suggest that altered interactions within the SHR ganglia reshape synaptic patterning as the animals develop and become hypertensive,

We generated primary cultures of postnatal WKY and SHR SNs grown in the absence or presence of SGCs to examine glial regulation of synaptic development. Cultures were grown for 14 days and stained for cholinergic synaptic markers (Fig. 8A). We confirmed that isolated SHR SNs had increased synaptic density compared to WKY SNs. Within each strain, however, addition of SGCs led to an increase in synaptic density (Fig. 8B left) and presynaptic marker expression (Fig. 8B right) compared to isolated SN cultures. These data are consistent with previous observations showing that SGCs contribute to the development of SN cholinergic networks (Enes et al., 2020). Given the higher baseline level of synaptic sites in the SHR SNs, we quantified the relative glial effect on synaptic density and found that SHR SGCs had a significantly smaller effect on the promotion of synaptic density and presynaptic marker expression than WKY SGCs (Fig. 8C). These data suggest that disrupted glial signaling in young SHR cultures limits synaptic development.

**Figure 8:**
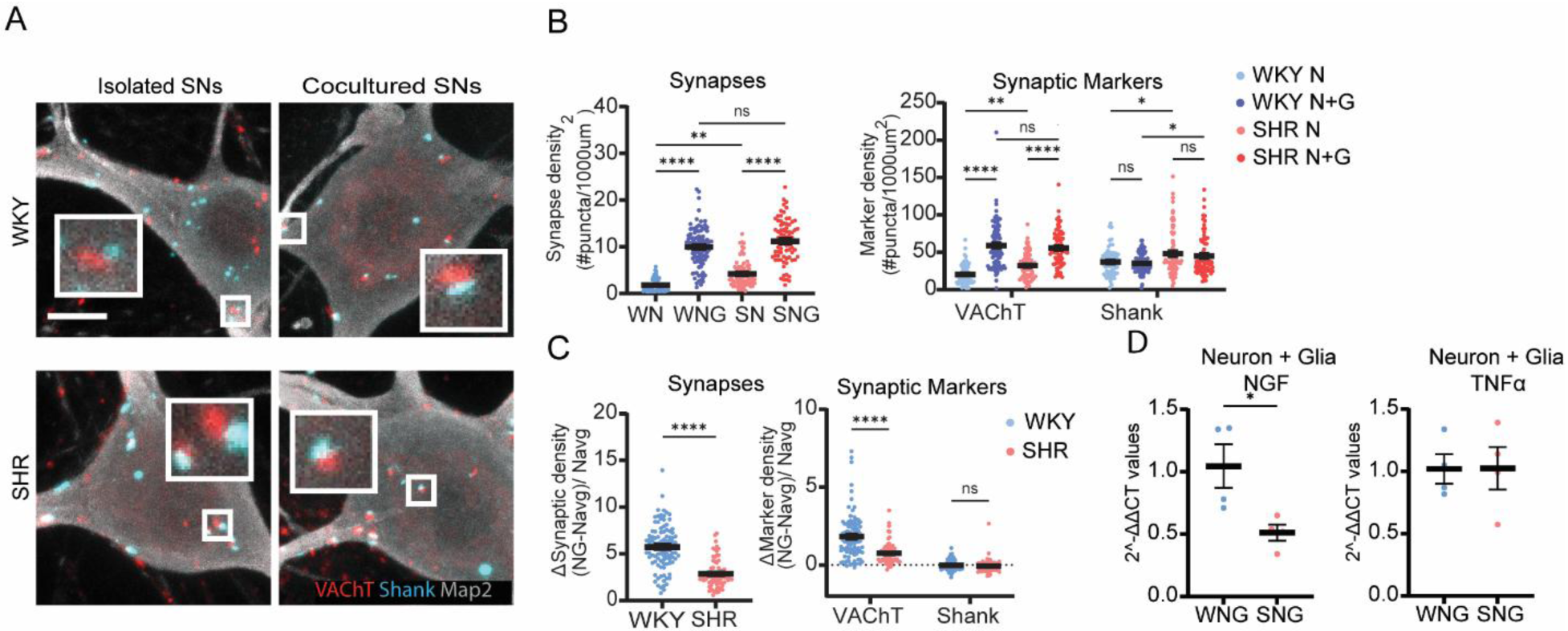
SGC-dependent regulation of synaptic density is altered in intrinsically hyperactive SHR cultures. **A,** Representative images of WKY and SHR SNs in isolated cultures (left column) and glial cocultures (right column) stained for VAChT, Shank, and MAP2. (Scale = 10 µm). **B,** Quantification of synaptic density (left) and individual pre- and postsynaptic markers (right) of WKY and SHR SNs cultured with SGCs (WNG and SNG) or in isolated cultures (WN and SN). (n=number of neurons; WN n=70, WNG n=100, SN n=76, SNG n=78. **Left**: WN vs WNG ****p<0.0001; WN vs SN ***p=0.005; SN vs SNG ****p<0.0001; WNG vs SNG n.s. **Right**: WN vs WNG VAChT ****p<0.0001, Shank n.s.; WN vs SN VAChT **p=0.0080, Shank *p=0.0131; SN vs SNG VAChT ****p<0.0001, Shank n.s.; WNG vs SNG VAChT n.s., Shank *p=0.0222). **C,** Quantification of the difference in synaptic density (left) and marker density (right) between SNs in cocultures vs isolated SNs cultures, (WN n=70, WNG n=100, SN n=76, SNG n=78. **Left**: Mann-Whitney test; WKY vs SHR ****p<0.0001. **Right**:2-way ANOVA with post-hoc Tukey’s multiple comparisons test; WKY vs SHR VAChT ****p<0.0001, Shank n.s.). **D,** Relative NGF (Left) and TNFα (Right) RNA expression in cocultures of SHR SNs and SGCs compared to WKY cocultures. (n=independent cultures; WKY n=4, SHR n=4. **Left:** WKY vs SHR NGF *p=0.0347. **Right:** WKY vs SHR TNFα n.s.).

We asked if the altered response to SHR SGCs in SN cocultures was associated with changes in the expression of glial synaptic regulatory molecules. We observed a decrease in the expression of NGF RNA in SHR compared to WKY cocultures (Fig. 8D left). In contrast, there was no significant difference in expression of TNFα RNA between the two strains (Fig. 8D right). This suggests that increased intrinsic activity in SHR neurons is associated with differential regulation of neurotrophic and pro-inflammatory factors in this disease model.

### Activity-dependent synaptic regulation is impaired in SHR neurons

We asked if activity-dependent synaptic compensation was disrupted in postnatal SHR SN cocultures by chemogenetically stimulating the SNs with CNO or vehicle for 48 hours (Fig. 9A). In contrast to the compensatory responses seen in WKY cocultures, CNO-treated SHR SNs had an increased synaptic density compared to the vehicle-treated SHR controls (Fig. 9B left). Surprisingly, we did not observe a change in presynaptic marker expression, however, CNO-treated SNs showed increased expression of post-synaptic markers (Fig. 9B right), consistent with altered plasticity mechanisms in pre-hypertensive SHRs.

**Figure 9:**
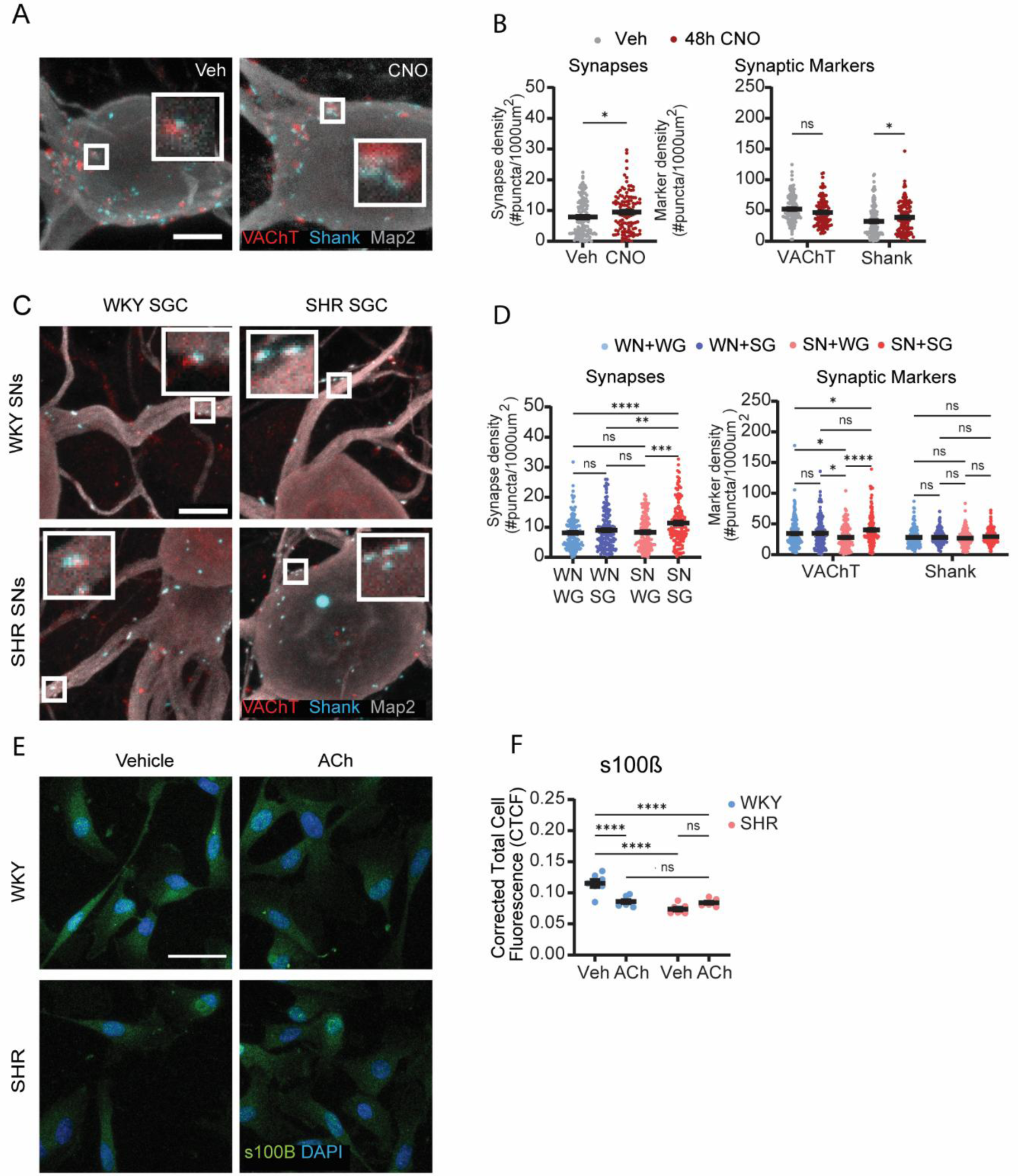
Loss of activity-dependent homeostatic regulation and altered glial properties in SHR cultures. **A,** Representative images of DREADD-expressing SHR SNs in cocultures treated with vehicle (Veh) or CNO and stained for VAChT, Shank, and MAP2 (Scale = 10 µm). **B,** Quantification of synaptic density (left) and individual pre- and postsynaptic markers (right) in DREADD-expressing SHR SNs in cocultures treated with Veh or CNO for 48 hours. (n=number of neurons; Veh n=108, 48h CNO n=96. **Left**: Veh vs. 48h CNO *p=0.0477. **Right**: Veh vs 48h CNO VAChT n.s., Shank n.s.). **C,** Representative images of strain-mixed cocultures: WKY SNs (top row) and SHR SNs (bottom row) cultured with WKY SGCs (left column) or SHR SGCs (right column) stained for VAChT, Shank, and MAP2. (Scale = 10 µm). **D,** Quantification of synaptic density (left) and individual pre- and postsynaptic markers (right) for strain-mixed cultures of WKY SNs cocultured with WKY (WN+WG) or SHR (WN+SG) SGCs and SHR SNs cocultured with WKY (SN+WG) or SHR (SN+SG) SGCs. (WN+WG n=158, WN+SG n=147, SN+WG n=135, SN+SG n=142. **Left**: WN+WG vs WN+SG, n.s.; WN+WG vs SN+WG n.s.; WN+WG vs SN+SG ****p <0.0001; WN+SG vs. SN+WG n.s.; WN+SG vs SN+SG **p <0.0031; SN+WG vs SN+SG ***p <0.0007. **Right**: WN+WG vs WN+SG VAChT n.s., Shank n.s.; WN+WG vs SN+WG VAChT *p=0.017 Shank n.s; WN+WG vs SN+SG VAChT *p=0.0309, Shank n.s.; WN+SG vs. SN+WG VAChT *p=0.0133, Shank n.s.; WN+SG vs. SN+SG VAChT n.s, Shank n.s.; SN+WG vs SN+SG VAChT ****p<0.0001, Shank n.s). **E,** Representative images of WKY and SHR SGCs treated with Veh or ACh for 24 hours and stained for s100β and DAPI. (Scale = 40 µm). **F,** Quantification of s100β corrected total cell fluorescence (CTCF) values for WKY and SHR SGCs treated with Veh or ACh. (n=independent culture dishes; WKY Veh n=7, WKY CNO n=7, SHR Veh n=7, SHR CNO n=7. WKY Veh vs. SHR Veh ****P<0.0001; WKY Veh Vs WKY Ach ****p<0.0001; WKY Veh vs. SHR ACh ***p<0.0001; SHR Veh vs. SHR ACh n.s.; WKY ACh vs. SHR ACh n.s.).

To further investigate the individual contributions of neurons and glia to disrupted synaptic regulation in the SHR cocultures we generated mixed cultures containing WKY SNs and SHR SGCs (WN-SG) or SHR SNs and WKY SGCs (SN-WG) and compared synaptic density to strain-matched cocultures (WN-WG and SN-SG) (Fig. 9C). Isolated SNs from SHR and WKY rats were cultured for 7 days before the addition of either SHR or WKY SGCs that had been grown separately for the same period. Synaptic density was measured after an additional 5 days (14 days total) of culture. Synaptic density was significantly higher in the SN-SG condition compared to all other coculture conditions (WN-WG, WN-SG, SN-WG) (Fig. 9D). Notably, SHR SNs do not show increased synaptic density in the presence of WKY SGCs. This suggests that the WKY SGCs possess compensatory activity that reduces the synaptic density of SHR SNs, and that this activity is lost in the SHR SGCs. Further, SHR SGCs do not drive increased synaptic density of WKY neurons, demonstrating an intrinsic hyperactivity profile of the SHR neurons that can be compensated by WKY, but not SHR SGCs. Together, these findings are consistent with alterations in both glial signaling and intrinsic neuronal properties in the SHR.

We further investigated glial changes associated with altered activity-dependent plasticity by examining the expression of s100β. S100β is a calcium-binding protein associated with a wide range of glial processes, including glial activation (Yardan et al., 2011), neurotrophic factor and cytokine release (Donato et al., 2009; Bianchi et al., 2010), and cortical plasticity (Nishiyama et al., 2002). We asked if s100β is regulated by cholinergic activation and whether there is differential regulation in WKY and SHR SGCs. We chronically treated isolated cultured WKY and SHR SGCs with ACh, stained for s100β and quantified the corrected total cell fluorescence (CTCF) of individual SGCs (Fig. 9E). Chronically stimulated WKY SGCs had significantly reduced s100β CTCF compared to vehicle-treated WKY SGCs (Fig. 9F). The baseline level of s100β was lower in SHR compared to WKY SGCs, with no significant change in response to ACh. This shows that a disrupted response to cholinergic transmission is associated with the loss of activity-dependent regulation in the SHR.

## Discussion

Mammalian peripheral functions such as blood pressure and cardiac output are normally maintained within a healthy range through a variety of homeostatic mechanisms including the regulation of autonomic neuronal activity (Ernsberger et al., 2021; Martinez-Sanchez et al., 2022; Habecker et al., 2025). Here we used chemogenetic and pharmacological methods to define an activity-dependent mechanism within the peripheral sympathetic nervous system that acts to homeostatically regulate cholinergic synaptic structural properties. This compensatory mechanism acts to bidirectionally constrain SN activity within a cultured cholinergic network. SGCs are necessary for compensatory responses to increased activity, as activity-dependent synaptic compensation was impaired when glial activation was disrupted or when SNs were stimulated in the absence of SGCs. Chemogenetic activation of the cultures also induced the downregulation of glial-expressed neurotrophic and cytokine genes associated with synapse formation, demonstrating that SGCs play an active role in activity-dependent regulation of sympathetic synapses. Finally, in a rat model of neurogenic hypertension marked by intrinsically elevated SN activity, we demonstrated altered activity-dependent plasticity and found a decrease in the capacity of SHR SGCs to restrain the strain-specific potentiation of SN synaptic density. Together these findings define the peripheral sympathetic system as a homeostatic regulatory site that influences SN output and suggests that loss of homeostatic structural plasticity could contribute to pathological disease progression.

Homeostatic synaptic plasticity in the CNS provides a compensatory mechanism that adjusts neuronal synaptic function and structural properties to constrain neuronal activity within a stable range (Wen and Turrigiano, 2024). Activity-dependent homeostatic plasticity in response to prolonged (24-48hr) changes in circuit activity triggers synaptic and firing rate regulation and has been described in cortical, hippocampal, and other brain circuits (Glazewski et al., 2017; Mendez et al., 2018; Wen and Turrigiano, 2021). Here we demonstrated that a structural form of synaptic compensation also exists in peripheral sympathetic circuits in response to chemogenetic and pharmacological manipulations of SN activity. While chronic stimulation induced compensatory synaptic reorganization, acute (2hr) chemogenetic stimulation did not alter synaptic density, even as presynaptic marker expression increased. This long-term compensatory mechanism may counter more acute forms of feedforward plasticity such as sympathetic short-term potentiation (Aileru et al., 2001) and gLTP (Brown and McAfee, 1982; Martínez et al., 2020).

Our findings indicate that activity-dependent homeostatic processes in the sympathetic system involve regulation of synaptic density, which decreased following prolonged periods of high activity and increased following sustained inhibition with TTX. Coordinate decreases in cFos expression following prolonged increased activity, as well as previous work demonstrating a link between increased synaptic density and increased synaptic transmission (Enes et al., 2020; Haburčák et al., 2022), further support a synaptic mechanism for homeostasis in the sympathetic circuit. This does not rule out, however, a possible role for regulation of intrinsic properties in sympathetic homeostatic plasticity. In the CNS, synaptic and firing rate changes can coordinately contribribute to homeostatic plasticity (Turrigiano, 2012; Wen and Turrigiano, 2024; Lu et al., 2025), although these processes have been shown to be uncoupled during development based on cricuit needs (Wen and Turrigiano, 2021). Changes in intrinsic excitability have been associated with homeostatic regulation in some SN preparations, but not in others. Activity-dependent up-regulation of SN excitability was reported in adult mice following spinal cord injuries that persistently reduced SN activity (Li et al., 2025). In contrast, angiotensin-II induced hypertensive mice that show increased cholinergic connections within the sympathetic ganglia did not have a compensatory decrease in intrinsic excitability (Li et al., 2024). Likewise, activity reductions in embryonic chick SNs showed little effect on intrinsic excitability (Ratliff et al., 2023). Further studies to measure SN firing rates following direct activity manipulations will be necessary to determine if changes in intrinsic excitability, in addition to our observed changes in synaptic properties, contribute to homeostatic adjustments in the sympathetic circuit.

We identified a requirement for a glial regulatory circuit for sympathetic activity-dependent plasticity. High neuronal activity triggered a glial response that signaled the SNs to reduce synaptic structures. This provides a direct homeostatic context for studies showing a role for SGCs in sympathetic synaptic function. Recently it has been shown that conditional depletion of sympathetic SGCs in a freely moving mouse enhanced sympathetic neuronal activity of the remaining SNs and functional output to the pupil, consistent with a compensatory response to reduced circuit activity (Mapps et al., 2022). In addition, acute chemogenetic activation of SGCs in mice resulted in an acute increase in heart rate, but a decrease in blood pressure during a more prolonged stimulation (Xie et al., 2017), again consistent with a homeostatic role for SGCs in the sympathetic circuit. Thus, our model of glial-dependent homeostatic regulation in a cultured sympathetic circuit can explain earlier studies involving *in vivo* manipulation of glial properties.

NGF is expressed by sympathetic SGCs (Enes et al., 2020), and is known to regulate synaptic development and function, increasing the expression of synaptic proteins (Lockhart et al., 2000), and synaptic activity via activation of the TrkA receptor (Luther et al., 2013). Here we showed chronic chemogenetic activation of cocultured SNs, and isolated SGCs, downregulated NGF expression and reduced synaptic density. Conversely, addition of exogenous NGF increased synapse number, consistent with a neurotrophic contribution to compensatory changes in this system. Decreased NGF levels have also been reported in cortical brain regions following increased neuronal activity associated with chronic stress paradigms (Ueyama et al., 1997; Scaccianoce et al., 2000), suggesting that neurotrophins may have a broader role in mediating homeostatic interactions.

In addition to NGF, direct stimulation of isolated SGCs also decreased TNFα expression. In the CNS, astrocytes provide a readily available pool of TNFα and depletion of TNFα from astrocytes disrupts homeostatic synaptic upregulation in hippocampal circuits (Heir et al., 2024). This work is consistent with a model in which activity-dependent downregulation of TNFα in sympathetic SGCs contributes to compensatory changes to decrease sympathetic synaptic density. It is interesting that in peripheral pain models TNFα acts as a pro-inflammatory factor that promotes increased sensory neuronal activity and pain behaviors (Tu et al., 2025). The idea that TNFα differentially regulates neuronal plasticity in disease models is consistent with our analysis of TNFα regulation in SHR cultures containing sympathetic SGCs and intrinisally hyperactive SNs. While expression of NGF and TNFα was downregulated by activity in WKY cultures, in the hyperactive SHR cultures this downregulation was seen for NGF, but not TNFα. Overall, this work suggests that neurotrophic and neuroimmune factors could act as glial-derived regulators of homeostatic plasticity and this regulation is disrupted in cells displaying pathological increases in actvity.

We identified cholinergic signaling as a trigger for activity-dependent changes in neurotrophin expression in SCGs. Chronic stimulation of SGCs with ACh downregulated NGF expression and, when neurons were present in the cultures, recapitulated the chemogenetic-induced compensatory regulation of synaptic density. Sympathetic SGCs have been reported to respond to acute ACh exposure via muscarinic receptors, resulting in increased intracellular calcium flux (Feldman-Goriachnik et al., 2018). In our system SGC activation was sufficient to drive synaptic compensation. This suggests that within the sympathetic circuit, SGCs respond to neurotransmitter spillover during chronic SN activation by decreasing levels of synapse promoting signals, resulting in compensatory loss of synapses (Fig. 6). In addition to NGF, we identified activity-dependent downregulation of TNFα and s100β, factors that have also been implicated in the modulation of synaptic properties (Nishiyama et al., 2002; Jaldeep et al., 2025). How these factors act and interact to modify synaptic plasticity will require further study.

Neurogenic hypertension is associated with increased SN activity that can be measured in humans (Esler and Kaye, 1998) and in rodent models (Doris, 2017; Davis et al., 2020; Li et al., 2024), including in the SHR rat and the angiotensin-induced hypertension mouse model. We showed that intrinsically hyperactive SHR SNs do not show synaptic compensation in the presence of SHR SGC, even though coculture with WKY SGCs eliminates excessive synapse formation. This suggests that the SHR SGCs do not detect and respond to the increased activity in the sympathetic circuit. The loss of ACh-dependent downregulation of s100β in the SHR glia provides further evidence of impairment in glial sensing of ongoing sympathetic activity. The idea that early neuronal hyperactivity is associated with loss of glial tracking of cholinergic signals may provide a context for understanding the observed increase in synaptic sites (Fig 7), associated with enhanced SN activity and hypertension in the adult SHR (Judy et al., 1976). Ongoing plasticity changes and persistent hyperactivity could contribute to the adult onset of hypertension and ultimately result in synaptic saturation, such as is seen in the limitation of gLTP in older adult SHRs (Alzoubi et al., 2010; Martínez et al., 2019). Together, this work suggests that a loss in homeostatic plasticity mechanisms contributes to unchecked feedforward plasticity in the SHR.

## Conflict of interest statement

The authors declare no competing financial interests

## Acknowledgments

This work was supported by National Institutes of Health (NIH) R21NS116316, the National Science Foundation (NSF) 1644666, the Baruchowitz Family Fellowship for Dysautonomia Research, and a grant from the W.M. Keck Foundation

## Notes

### Competing Interest Statement

The authors have declared no competing interest.

### Summary of Updates

Figure 7-9 revised. Fixed grammar and formatting on some sections. Adjusted some language and how we frame the story.

